# Emerging Resistance to Novel β-Lactam β-Lactamase Inhibitor Combinations in *Klebsiella pneumoniae* bearing KPC Variants

**DOI:** 10.1101/2025.04.14.648765

**Authors:** Mase Hamza, German M. Traglia, Lucia Maccari, Sonia Gomez, Maria Belen Sanz, Usman Akhtar, Vyanka Mezcord, Jenny Escalante, Alejandra Corso, Cecilia Rodriguez, Christopher R. Bethel, Gauri G. Rao, Marcelo E. Tolmasky, David Paterson, Robert A. Bonomo, Fernando Pasteran, Maria Soledad Ramirez

## Abstract

**Background:** *Klebsiella pneumoniae* carbapenemase (KPC) variants, predominantly KPC-2 and KPC-3, are significant global resistance mechanisms. KPC-2 and KPC-3 confer resistance to a broad range of β-lactams, including carbapenems, while remaining susceptible to ceftazidime-avibactam (CZA). Recently, new KPC variants have developed resistance to CZA through mutations, insertions, or deletions in regions such as the Ω-loop, 240-loop (237–243 aa), and 270-loop (266–275 aa). This study aimed to investigate the collateral resistance to cefiderocol (FDC) and cefepime/zidebactam (FPZ) among isolates with these mutations.

**Methods:** Fifteen clinical isolates of KPC-producing *Klebsiella spp*. were analyzed, representing 15 distinct variants. Antimicrobial susceptibility testing determined the MICs for CZA, carbapenems, FDC, FPZ, and other antibiotics. Synergy between CZA and FDC was assessed. Whole-genome sequencing (WGS) was used to identify mutations contributing to resistance.

**Results:** CZA resistance was confirmed in 12 of the 15 KPC variants. Collateral resistance to FDC was observed in eight isolates, with five exhibiting spontaneous resistant subpopulations. Six FDC-resistant strains had mutations in the 270-loop (266–275 aa). Collateral resistance to FPZ was seen in three KPC variants, especially those with mutations in the 270-loop (266–275 aa), though many Ω-loop and 240-loop (237–243 aa) mutants remained susceptible. WGS of FDC-resistant subpopulations revealed additional mutations in *ompC, rpoC, dksA*, and *cirA*.

**Conclusions:** This study demonstrates that emerging KPC variants showing resistance to CZA also exhibit resistance to FDC, with collateral resistance to FPZ observed to a lesser extent. Identifying mutations in *bla*_KPC_, *cirA*, and other genes is important to understand resistance mechanisms for effective therapies.

## INTRODUCTION

*Klebsiella pneumoniae* carbapenemase (KPC) variants, specifically KPC-2 and KPC-3, have gained significant global prevalence, particularly in *K. pneumoniae* isolates [1-3]. These β-lactamases provide resistance to most β-lactams, including carbapenems, but remain susceptible to newer β-lactam/ β-lactamase inhibitors like ceftazidime-avibactam (CZA). However, since the introduction of CZA, numerous KPC variants resistant to this combination have emerged [2, 3]. In these KPC variants, mutations, insertions, and/or deletions have been identified in distinct regions of KPC β-lactamase. Mutational “hot spots” associated with resistance to CZA are located in specific regions: (i) the Ω-loop (residues 164 to 179, which border the lower part of the catalytic pocket), (ii) the 240-loop (amino acids 237 to 243. adjacent to the conserved KTG motif and defining the right side of the active site), and (iii) the 270-loop (amino acids 266 to 275, positioned further from the active site between β strand 5 and the α11 helix) [3-6]. The emergence of ceftazidime-avibactam resistance in KPC-producing *K. pneumoniae* has been associated with mutations in these critical structural regions. Recent studies have described novel KPC variants, such as KPC-189 and KPC-197, which confer resistance to CZA through modifications in these sites [7, 8]. Additionally, the KPC-74 variant has been identified as a CZA-resistant enzyme that emerged during treatment, highlighting the ongoing evolution of KPC enzymes under selective pressure[9].

Understanding the effects of these substitutions is essential for tracking the evolution and spread of KPC variants resistant to CZA and its collateral resistance to other antibiotics, such as cefiderocol (FDC) and cefepime/zidebactam (FPZ).

Published literature have reported the occurrence of collateral resistance to FDC in few KPC variants, including KPC-31, KPC-33, KPC-62, and novel variants like KPC-109 and KPC-203 (Table S1). The concept of “collateral resistance” has been described in the literature as the unintended resistance to one antibiotic due to selective pressure exerted by another, even when they do not share the same direct target [10, 11]. Specifically, mutations in *bla*_KPC_ associated with CZA resistance have been linked to structural changes in the enzyme that impact its interaction with other β-lactams, including FDC [12]. Similarly, porin alterations— often secondary to β-lactamase evolution—can restrict FDC uptake, further supporting the collateral resistance phenomenon [13]. Most of these variants exhibiting FDC collateral resistance have substitutions in the Ω-loop region, significantly affecting the hydrolysis of both antibiotics and contributing to FDC collateral resistance. Notably, KPC-109, a variant of KPC-3, was identified in a clinical isolate (NE368) with a six-amino acid insertion in the 270-loop region, mediating resistance to CZA and FDC[14].

To our knowledge, collateral resistance to FPZ and cefepime/taniborbactam in clinical isolates has not been reported in the literature. The aim of our present study is to describe the occurrence of collateral resistance to FDC and FPZ in *Klebsiella* clinical isolates harboring various KPC variants (n=15) in the three main specific regions.

## MATERIALS and METHODS

### Bacterial strains

A total of 37 out of 175 *Klebsiella* spp. clinical isolates harbored a KPC variant distinct from KPC-2 or KPC-3. A representative isolate from each variant was selected for this study. A total of fifteen selected KPC-producing *Klebsiella* spp. clinical isolates, not epidemiologically related and collected from various regions in Argentina over the span of four years (2020-2023), were included (Table S2). Among these isolates, 15 KPC variants were included (Table 1 and Figure 1). The *K. pneumoniae* ATCC reference strains BAA1705 (KPC-2 producer) and 13883 (susceptible) were also tested as reference.

**Table 1.**
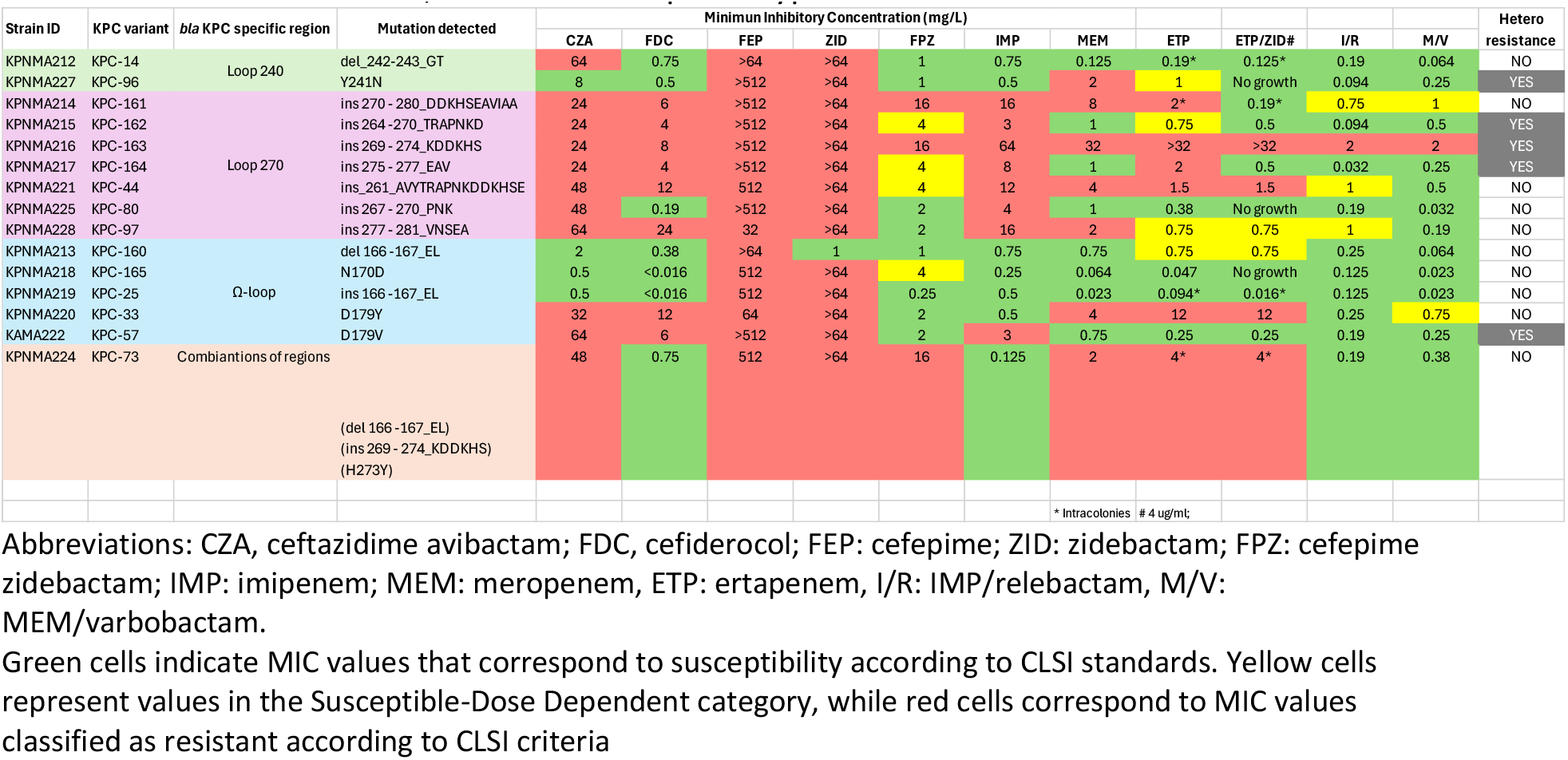
KPC variants strains, molecular and phenotypic characteristics.

**Figure 1:**
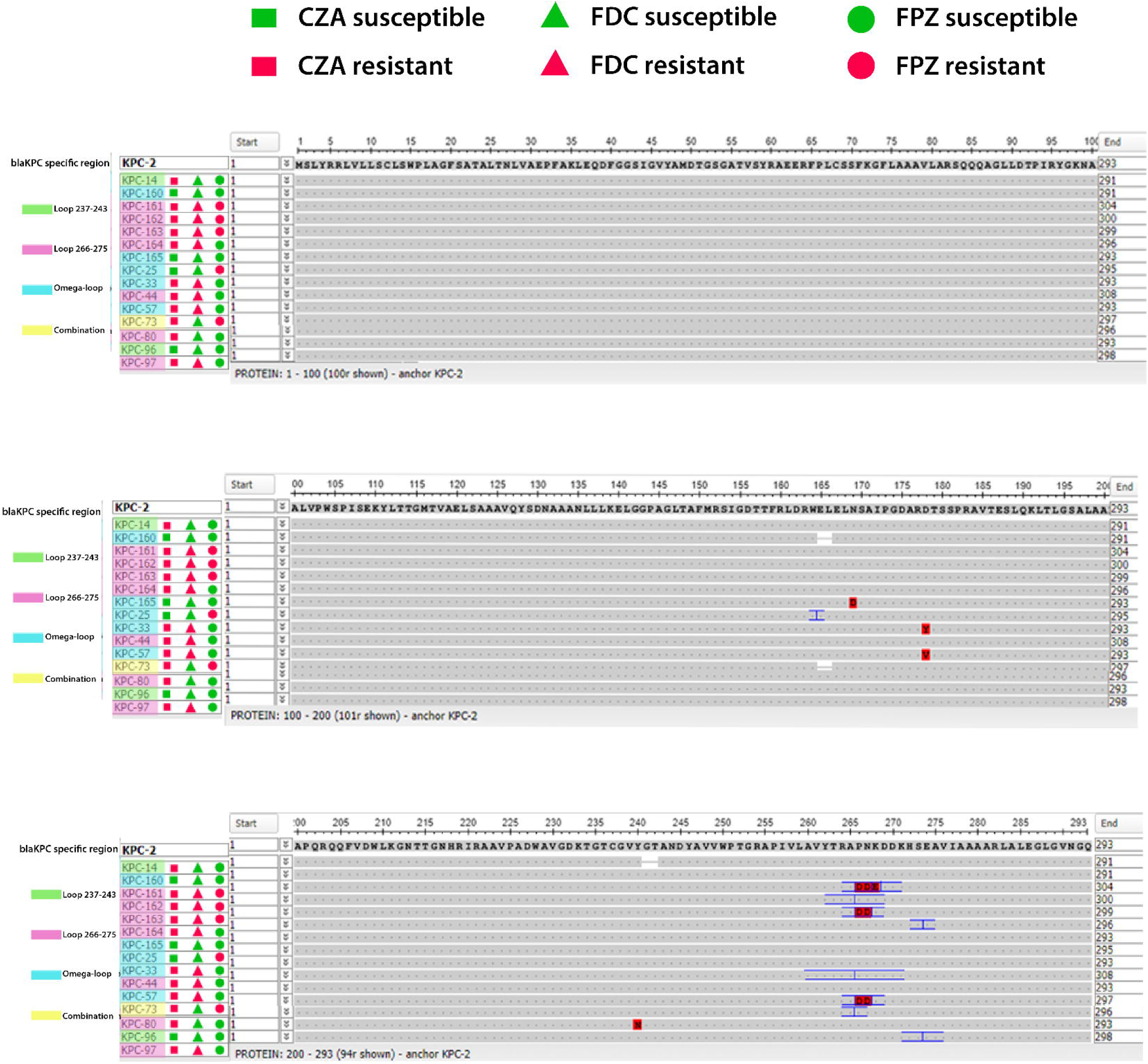
KPC variants aligned and compared against KPC-2. Protein sequences were retrieved from the CARD database and β-Lactamase Database. Aligned with MUSCLE, visualized by NCBI MSA viewer.

### *Antimicrobial susceptibility testing* (AST)

The minimum inhibitory concentrations (MICs) for CZA, meropenem (MEM), imipenem (IMP), cefepime (FEP), zidebactam (ZID), FPZ, ertapenem (ETP), ertapenem/zidebactam (ETP/ZID*), imipenem/relebactam (I/R), and meropenem/varbobactam (M/V) were determined using broth microdilution methods and/or commercial E-strips (Liofilchem S.r.l., Roseto degli Abruzzi, Italy) in accordance with the Clinical and Laboratory Standards Institute (CLSI) guidelines. For testing FDC susceptibility, commercial E-strips (Liofilchem S.r.l., Roseto degli Abruzzi, Italy) and broth microdilution with iron-depleted cation-adjusted Mueller-Hinton medium as the reference method were used. All procedures were carried out in accordance with the manufacturer’s instructions and met the standards of CLSI [15] and the European Committee on Antimicrobial Susceptibility Testing (EUCAST) (https://www.eucast.org/clinical_breakpoints).

Resistant subpopulations within the inhibition ellipse zones of FDC were selected for further studies and whole genome sequence analysis. Stocks of the FDC resistant subpopulations were stored at -80°C and the stability of the FDC resistance was determined.

For FPZ categorization, the CLSI FEP breakpoint was used. Synergy between CZA and FDC was perform in resistant and heteroresistant strains using MTS™ Synergy Application System (Liofilchem S.r.l., Roseto degli Abruzzi, Italy). Synergy was evaluated using the fractional inhibition concentration index (FICI) as previously described [16]. Quality control strains, such as *Escherichia coli* ATCC 25922 and the *K. pneumoniae* ATCC 13883 strains were included in the experiments. Each strain was tested at least in duplicates.

### Whole genome sequencing analysis

The genomic DNA extraction of parental strain and intra halo colonies was performed using Wizard Promega kit (Promega, Madison, WI, USA). The Genomic sequencing was done using NovaSeq X Plus, producing 2×151 bp paired-end reads. To ensure sequence quality, FASTQC software analysis (https://www.bioinformatics.babraham.ac.uk/projects/fastqc/) was performed, followed by trimming and filtering using Trimmomatic software (version: 0.40, ILLUMINACLIP: TrueSeq3-PE.fa.2:30:10; LEADING:3; TRAILING:3; SLIDINGWINDOW: 4:15; MINLEN:36) [17]. De novo sequence assembly was conducted using SPAdes (version: 3.15.4, default parameters) [18] and the quality was subsequently evaluated using QUAST (version: 5.2.0) [19]. Genome annotation was performed through PROKKA (version 1.14.5)[20], while variant calling was carried out using the *breseq* and *gdtools* software packages (version: 0.38.1, consensus mode, default parameters) [21]. Recombination regions were identified and removed using Gubbins software (version: 3.3.0, default parameters) [22]. Plasmid identification was performed using PlasmidFinder v2.1 [23]. The copy number gene (*bla*_KPC_) was assessed using CCNE tool [24]. The Whole Genome Shotgun project of KPNMA215, KPNMA216 and KAMA222 strains has been deposited in GenBank under accession numbers JBIUGI000000000, JBIUGH000000000, and JBIUGG000000000respectively. In addition, the fastq files, quality analysis files, assemblies’ sequences and genome annotation files of wild-type and IHC were uploaded in Zenodo repository: https://zenodo.org/records/15122353(last accessed April 1, 2025).

## RESULTS AND DISCUSSION

### Description of CZA resistance and collateral resistance to FDC in *Klebsiella* KPC variants strains

The CZA MICs for 15 KPC variants were determined, confirming resistance in 12 of them (Table 1). Interestingly, three variants with mutations in the Ω-loop—KPC-25, KPC-160, and KPC-165— remained susceptible to CZA, despite alterations in this critical region (Table 1). Prior publications of these specific KPC variants have been identified in the literature.

We next studied the collateral resistance to FDC, and we observed resistance in eight of the 15 isolates, using EUCAST breakpoint (BP) guidelines. Among the tested strains, resistant colonies with the gradient strip inhibition zone (FDC resistant subpopulations) were also observed in five strains including one that was categorized as FDC susceptible (Table 1). Six of the FDC resistant strains harbor substitutions in the 266–275 loop (Table 1). Previous reports have identified cross resistance to FDC when substitutions in the Ω-loop are present (Table S1) [3]. In addition, these studies highlight the emergence of FDC resistance in high-risk clones such as the ST307 lineage, which has been responsible for outbreaks in hospital settings, particularly in ICUs [25, 26]. KPC-62, identified in ST307 isolates, displayed resistance to both CZA and FDC, driven by the L169Q mutation within the *bla*_KPC_ gene. The novel KPC-203 variant found in Italy adds further complexity to the resistance landscape, as it exhibits cross resistance to CZA and FDC due to significant modifications at key positions, including a deletion in the Ω-loop and an insertion in the 260 amino acid position [27]. The newly identified KPC-216 shows cross-resistance to both CZA and FDC. The KPC-216 variant, characterized by a lysine insertion at position 170 in the Ω-loop, was isolated from a *K. pneumoniae* ST101 strain and demonstrated resistance to both CZA and FDC [28]. In addition, KPC-109, a variant of KPC-3, was identified in the clinical isolate, featuring a six-amino acid insertion in the 270-loop region, which conferred resistance to both CZA and FDC [14]. Reports with mutations such as the D179Y substitution found in both KPC-31 and KPC-33, and observed in this work, illustrate the challenge of treating infections with these variants, as they lead to not only CZA resistance but also a potential for cross-resistance with FDC [29-32].

We tested the synergy between CZA and FDC in all strains exhibiting resistance or heteroresistance to both antibiotics. Synergy was observed in four strains (Table 2). Particularly, synergy was seen in the strains exhibiting intracolonies for FDC (Table 1 and Table 2). Evaluating the synergy between FDC and CZA, a clinically available fixed combination, addresses the need to explore relevant therapeutic options. While avibactam has previously demonstrated synergy with FDC, it is not available as a stand-alone agent for clinical use. In contrast, CZA is a formulation used in clinical practice that contains both ceftazidime and avibactam. Therefore, assessing the synergy between CZA and FDC allows for a more translationally relevant interpretation of potential combination therapies.

**Table 2.** Synergy between CZA and FDC for FDC resistant/hetero-resistant KPC variant strains.

| MTS synergy |  |  |  |  |
| --- | --- | --- | --- | --- |
| Strains | MIC CZA(FDC) | MIC FDC(CZA) | FICI | Synergy |
| KPNMA214 | 6 | 2 | 0.58 | Additive |
| KPNMA215 | 4 | 0.5 | 0.29 | Synergy |
| KPNMA216 | 6 | 1 | 0.38 | Synergy |
| KPNMA217 | 0.75 | 0.38 | 0.13 | Synergy |
| KPNMA220 | 16 | 4 | 0.83 | Additive |
| KPNMA221 | 64 | 4 | 1.67 | Additive |
| KAMA222 | 4 | 0.094 | 0.08 | Synergy |
| KPNMA227 | 3 | 0.19 | 0.76 | Additive |
| KPNMA228 | 32 | 8 | 0.83 | Additive |

FDC: cefiderocol, FICI: Fractional Inhibitory concentration index.
Synergy was defined as FICI ≤ 0.5, Additive effect as FICI between 0.5 and 1.

### Impact of KPC variants in cefepime/zidebactam susceptibility

Firstly, we observed that all 15 KPC variants were resistant to cefepime (FEP), and all but one were resistant to zidebactam (ZID) (Table 1). When testing FPZ, three of the KPC variants displayed resistance (Table 1). Notably, most strains harboring mutations in the Ω-loop and the 240-loop remained susceptible to FPZ.

Only one report on FPZ has been published, where the authors evaluated the antimicrobial susceptibility of a KPC-33 variant, which exhibited susceptibility to FPZ (Shi Qingyu, 2020, DOI: 10.1093/CID/CIAA1521). Another report evaluated cefepime/taniborbactam, showing decreased susceptibility in laboratory-constructed strains with substitutions in both the Ω–loop and the 240-loop, particularly when combined with porin defects (mutations in OmpK35 and OmpK36) [33]. Our study is the first to describe collateral resistance to FPZ in KPC variants.

When compared with KPC-producing clinical isolates from the same period that lacked KPC mutations, variants exhibiting resistance to CZA were significantly more likely to be resistant to FPZ and FDC and more susceptible to carbapenem (Fisher’s exact test, p < 0.05) [34]. While based on limited samples, this comparison and analysis fit for normal distribution and open new venues for future studies with larger cohorts.

As amino acid changed in the MarR, PBP-2, PBP-3, OmpK35, and OmpK36 proteins, or their respective promoters, may contribute to the FDC and FPZ resistance phenotypes [35, 36]. We analyzed the sequences of the three KPC-producing strains with FPZ resistance. A comparative analysis of KPNAMA215 and KPNAMA216 against the FDC-susceptible ATCC 13883 strain revealed no amino acid substitutions in PBP-2 and PBP-3. However, in OmpK35, only KPNAMA216 exhibited an A8T substitution, while multiple amino acid changes were observed in OmpK36 in both KPNAMA215 and KPNAMA216 (Fig. S1).

The comparison of the KAMA222 genome to *K. aerogenes* FDAARGOS 1442 (used as the reference genome) identified two amino acid substitutions in PBP-2 (G107D and A354D) (Fig. S2), as well as multiple changes in OmpK36, consistent with the findings in KPNAMA215 and KPNAMA216 (Fig. S1 and Fig. S2). In MarR, an S82G amino acid substitution was detected in KPNAMA215 and KPNAMA216, whereas KAMA222 showed no amino acid changes in this protein (Fig. S2). The presence of multiple amino acid substitutions in OmpK36 across the analyzed genomes complicates the assessment of their specific contribution to FDC and FPZ resistance. However, we cannot rule out the possibility that these alterations play a role in the observed resistance. Further analysis involving a larger number of strains and their respective isogenic strain with not mutation on the porins is necessary to draw definitive conclusions. In conclusion, while previous reports have highlighted susceptibility to FPZ in specific KPC variants, such as KPC-33, and documented reduced susceptibility in other FEP combinations like FEP/taniborbactam under certain genetic conditions, our study is the first to describe collateral resistance to FPZ in multiple KPC variants. This finding underscores the complexity of resistance mechanisms in KPC-producing strains, particularly those involving mutations in the 270-loop (266-275 aa).

### Collateral resistance to ertapenem/zidebactam (ETP/ZID) KPC variants

Among the 15 KPC variant isolates tested in this study, six were resistant to ertapenem (ETP) (Table 1). When evaluating the ETP/ZID combination, susceptibility was restored in only two isolates (Table 1). Most of the ETP-resistant strains harbored mutations in the 270-loop (266-275 aa), which may contribute to their resistance phenotype (Table 1).

Notably, one isolate (KPNMA216) exhibited the highest MIC (>32 mg/L) to ETP and ETP/ZID, and this strain possessed alterations in both OmpK35 and OmpK36 and the 270-loop (266-275 aa) mutations on the KPC (Table 1 and Fig. S1). The presence of porins mutations suggests a potential role in limiting β-lactam uptake and enhancing resistance to β-lactam/β-lactamase inhibitor combinations.

Although all the isolates analyzed in this study come from Argentina, the resistance mechanisms identified, in particular the mutations in *bla*_KPC_ associated with resistance to CZA and its impact on FDC and FPZ, have been reported in various regions of the world, including in high-risk clones such as ST258 and ST307 [7, 25, 37, 38]. Our findings agree with these previous studies, suggesting that the described mechanisms can be extrapolated beyond the local context.

To date, there have been no prior reports of collateral resistance to FDC and ETP/ZID in KPC-producing isolates [39-41]. This study showed the occurrence of this phenomenon for the first time and underscores the critical need for ongoing comprehensive surveillance to monitor emerging resistance patterns in multidrug-resistant *Klebsiella*.

### Collateral susceptibility to carbapenems in *K. pneumoniae* KPC variants

We tested the MICs of imipenem (IMP) and meropenem (MEM) in the 15 KPC variants, and we observed in 12 strains collateral susceptibility to IMP (n=1), MEM (n=5), and both (n=6) (Table 1). MEM exhibits lower MICs compared to imipenem, indicating that it is generally less affected by resistance mechanisms, including *bla*_KPC_ variants. Cross resistance to carbapenems was seen in strains harboring KPC variants with substitutions in the 270-loop (266-275 aa) (Table 1). The mutations in the 270-loop region can alter the local flexibility and conformation of the active site, influencing substrate binding and catalytic efficiency, particularly in class A β-lactamases [42-44].

Previous investigations have suggested that certain substitutions in KPC variants, particularly those in the Ω-loop region, may reduce the enzyme’s carbapenemase activity, thereby restoring susceptibility to carbapenems [3, 4, 45, 46]. For instance, variants such as KPC-31 and KPC-33, which harbor the D179Y substitution, have been shown to confer resistance to CZA while exhibiting collateral susceptibility to carbapenems. No isolates with mutations in 270-loop (266-275 aa) showed collateral susceptibility to imipenem. Although our study did not include kinetic validation, previous literature has demonstrated the functional relevance of these mutations through mechanistic approaches, showing that substitutions, insertions or deletions can contribute to CZA resistance by increasing affinity for CAZ and reducing susceptibility to AVI, ultimately affecting enzyme stability and broadening its spectrum of activity [47] [3, 9]. In addition, we tested imipenem/relebactam (I/R) and meropenem/varbobactam (M/V) and we observed that vaborbactam reduced meropenem MICs by more than 2 log_2_-fold in 12 strains, whereas relebactam achieved the same reduction in 8 out of 15 strains. (Table 1).

Our work and previous findings highlight the complex interplay between β-lactamase mutations and antibiotic efficacy, where changes that drive resistance to one class of antibiotics may inadvertently restore susceptibility to others, such as carbapenems. More research is needed to further explore this phenomenon across different KPC variants.

### WGS analysis of FDC spontaneous emergent resistant cells

Spontaneous hetero-resistant colonies from three KPC variants strains were chosen for further analysis. The MIC for FDC for these resistant subpopulations showed a 2-fold increase for KPNMA215 and KPNMA216 and a 3-fold increase KPNMA222 (Table 3).

**Table 3.**
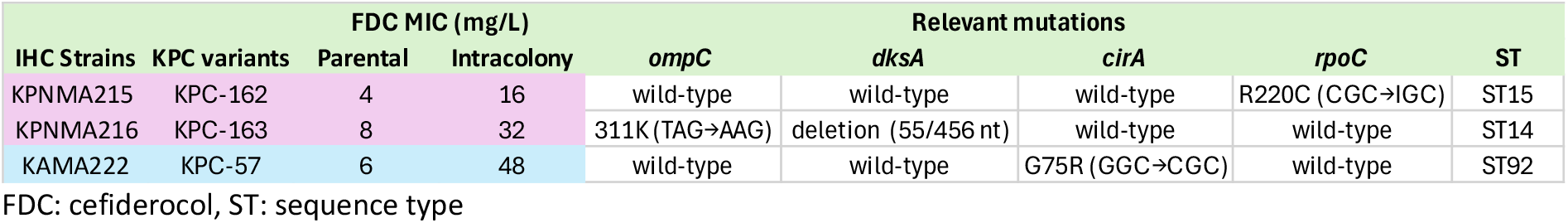
Parental and resistant subpopulations FDC MIC and ST, and relevant mutations associated with antibiotic resistance.

The WGS was performed and subsequent genome analyses of *K. pneumoniae* strains KPNMA215 and KPNMA216, along with *K. aerogenes* strain KAMA222 and their FDC-resistant subpopulations, revealed distinct sequence types (STs) and species classifications. KPNMA215 was classified as ST15, KPNMA216 as ST14, and KAMA222 as ST92. Common antimicrobial resistance (AMR) genes were found across all strains, including *bla*_TEM_, *bla*_SHV_, *oqxA, oqxB, fosA6, tet(A)*, and *tet(D)*, as well as aminoglycoside-modifying enzymes such as *aac(6’)-Ib-cr* and *aph(6)-Id* (Table S3). The presence of plasmids was identified in all three sequenced genomes. All three genomes harbor plasmids belonging to the incompatibility group IncL/M. Additionally, an IncFII-type plasmid was found in KPNAMA215 and KPNAMA222. An IncFIB-type plasmid was identified in the genomes of KPNAMA215 and KPNAMA216. Furthermore, KPNAMA215 also carries a plasmid belonging to the incompatibility group IncR. No resistance genes were identified on the plasmids.

In the mutational analysis of the FDC-resistant subpopulation IHC215, a mutation was detected in the *ISSod9* transposase, which plays a role in the movement of genetic elements and may facilitate the horizontal transfer of resistance genes. Additionally, a mutation was found in the *rpoC* gene, which encodes the β-subunit of DNA-directed RNA polymerase.

Mutations in *rpoC* have been associated with transcriptional accuracy and stress adaptation, which may contribute to bacterial survival under antibiotic pressure by modulating global gene expression. The mutation identified in *rpoC* in IHC215 may alter RNA polymerase function and global gene expression patterns, ultimately reducing susceptibility to FDC (Table 3 and Table S4).

For KPNMA216 FDC-resistant subpopulation IHC216, mutations were detected in the *ompC* gene. This porin facilitates passive diffusion of β-lactams into the periplasmic space, and its alteration likely reduces permeability, limiting antibiotic entry and contributing to resistance. Similar resistance mechanisms have been observed in clinical isolates involving OmpK35 and OmpK36 porins in *K. pneumoniae* [27, 33, 48]. The presence of the *ompC* mutation may contribute to the reduced susceptibility to antibiotics [33]. A mutation in *dksA* gene, which encodes an RNA polymerase-binding protein, was also observed, suggesting its role in regulating gene expression under stress, particularly during antibiotic exposure (Table 3 and Table S4) [49]. Specifically, a deletion of the first 93 nucleotides was identified in the *dksA* gene, potentially leading to a loss-of-function mutation. This truncation may impair the bacterial stress response and further modulate the expression of resistance or persistence-related genes (Table S4). Lastly, in the IHC222 subpopulation, we identified a typical mutation in the *cirA* gene, a gene frequently prone to mutations and previously linked to FDC resistance (Table 3 and Table S4). *cirA* encodes a TonB-dependent outer membrane receptor responsible for ferric-siderophore uptake. Although the mutation identified here has not been previously reported, its location and nature suggest functional inactivation. The novelty of this mutation highlights the genetic flexibility of *cirA* and its role as a resistance determinant. A previous case study of a hypervirulent *K. pneumoniae* strain co-producing KPC-2 and SHV-12 identified a truncation in *cirA*, leading to FDC resistance [50]. Similarly, other studies have shown that *cirA* mutations, when combined with the production of NDM-5 carbapenemase, result in even higher levels of FDC resistance [26, 51-54]. This synergy between *cirA* inactivation and β-lactamase activity underscores the complex interplay of resistance mechanisms. Previous findings and our current work emphasize the need to monitor *cirA* mutations as a significant marker of FDC resistance in clinical settings, particularly in strains producing KPC-2, and KPC variants, as well as NDM-5.

The copy number of the *bla*_KPC_ gene and the associated increase in carbapenems, CZA, and β-lactamase/inhibitors resistance have been previously reported [35, 48, 55]. In this study, the copy number of the *bla*_KPC_ gene in both WT and IHC colonies of strains KPNAMA215, KPNAMA216, and KAMA222 was evaluated using whole genome sequencing data and the CCNE tool. An increased *bla*_KPC_ copy number was observed in the IHC colonies of KPNAMA215 (1.52-fold changes) and KAMA222 (1.48-fold changes) compared to their respective WT colonies. In contrast, no difference in the *bla*_KPC_ copy number was detected in the IHC colonies of KPNAMA216.

Together, our findings indicate that mutations in *cirA, ompC, rpoC*, and *dksA* play complementary roles in reducing susceptibility to FDC by affecting membrane permeability, drug uptake, and transcriptional regulation under stress conditions. These mechanisms highlight the multifactorial nature of resistance and the importance of monitoring such mutations in clinical surveillance.

A limitation of our study is that, although we identified mutations potentially associated with resistance, such as in the *ompC* and *rpoC* genes, we did not experimentally validate their functional impact using gene knockout or complementation assays. Although these experiments would provide functional confirmation, our primary objective was to identify mutational events potentially related to FDC and CZA resistance in clinical isolates with KPC variants, employing genomic surveillance and comparative sequence analysis. Nevertheless, our findings underscore the need for future research to determine the functional impact of these mutations on the resistance phenotype.

## Conclusion

This study highlights the complex genetic landscape of KPC variants and the emergence of collateral resistance to both FDC, FPZ and ETP/ZID. Notably, several variants, particularly those harboring mutations in critical regions such as the Ω-loop and the 270-loop, contributed to high levels of resistance to CZA and FDC. Collateral resistance to FPZ was observed to a lesser extent, specifically among variants with mutations in the 270-loop. The detection of collateral resistance to both antibiotics in strains that had not been previously exposed to these drugs is particularly concerning, as it suggests the potential for resistance to develop even without selective pressure from these antibiotics. Additionally, our findings revealed spontaneous mutations in FDC-resistant subpopulations, including previously known mutations in the *cirA* gene as well as novel mutations not previously associated with FDC resistance. This underscores the importance of continuous monitoring of KPC variants, especially in clinical settings, to prevent further spread of resistance and to develop more effective treatment strategies as KPC allelic variants may influence susceptibility profiles, particularly regarding their response to novel β-lactam agents. Understanding the molecular mechanisms driving resistance will be crucial for managing and controlling the spread of these multidrug-resistant pathogens.

## Author Contributions

GMT, GGR, MET, RAB, FP, and MSR conceived the study and designed the experiments. MH, GMT, LM, SG, MBS, CR, UA, VK, JE, FP, and MSR performed the experiments and genomics and bioinformatics analyses. GMT, FP, CRB, GGR, DP, MET, RAB, and MSR analyzed the data and interpreted the results. GMT, FP, MET, and MSR contributed reagents/materials/analysis tools. GMT, FP, GGR, CRB, AC, MET, RAB, and MSR wrote and revised the manuscript. All authors read and approved the final manuscript.

## Supplementary Material

**Figure S1**.

Comparison of the OmpK35, OmpK36, and MarR amino acid sequences between KPNAMA215, KPNAMA216, and *Klebsiella pneumoniae* ATCC 13883, using the latter as the reference genome from the NCBI Genome database.

**Figure S2**.

Comparison of the OmpK36 and PBP-2 amino acid sequences between KAMA222 and *Klebsiella aerogenes* FDAARGOS 1445, using the latter as the reference genome from the NCBI Genome database.

**Table S1:** Reported cases in the literature highlight the occurrence of collateral resistance to FDC in KPC variants

**Table S2:** Demographic data and characteristics of the fifteen *Klebsiella* spp. strains included in this manuscript.

**Table S3:** Antimicrobial resistance genes present in KPNMA215, KPNMA216 and KAMA222.

**Table S4:** Mutational analysis of KPNMA215, KPNMA 216, and KAMA222 IHCs compared to the corresponding parental strain.

