## Supplementary material for "Emerging Resistance to Novel β-Lactam β-Lactamase Inhibitor Combinations in *Klebsiella pneumoniae* bearing KPC Variants": Suplemental Material: Table S1 and S2 NEW.pdf

**Table S1.** Reported cases in the literature highlight the occurrence of collateral resistance to FDC in KPC variants.

| *bla*KPC Variants | Clinical or Laboratory Isolates | Region of Protein Mutated | CZA MIC (mg/L) | FDC MIC (mg/L) | DOI |
|---|---|---|---|---|---|
| KPC-109 | *K. pneumoniae* NE368 | (ins 268 - 273_NKDDKY) (H273Y) | >256 | >8 | DOI |
| KPC-33 | Laboratory strain (KPC-2 mutant) | D179Y | 256 | 4 | DOI |
| KPC-62 | *K. pneumoniae* (ST307, ICU) | L169Q | ≥16/4 | ≥4 | DOI |
| KPC-203 | *K. pneumoniae* ST101 | (del 166-167_EL) (ins 260-268_LAVYTRAPM) | Cross-resistant | Cross-resistant | DOI |
| KPC-31 | *K. pneumoniae* ST307 | D179Y | >16 | Cross-resistant | DOI, DOI |
| KPC-50 | Clinical isolates | (ins 275 – 277_EAV) (H273Y) | Resistant | Resistant | DOI |
| KPC-33 and KPC-2 | Clinical isolates (KPJCL-3 and KPJCL-4) | D179Y mutation | >32 | Reduced susceptibility | DOI |

**Table S2.** Demographic data and characteristics of the fifteen *Klebsiella* spp. strains included in this manuscript.

| Strain ID | Species | KPC variant | Sex | Age | Data of isolation | Source of isolation | Other relevant *bla* | Hospital |
|---|---|---|---|---|---|---|---|---|
| KPNMA212 | *K. pneumoniae* | KPC-14 | M | 62 | 7/2/21 | urine | CTX-M-15 | H1 |
| KPNMA227 | *K. pneumoniae* | KPC-96 | F | NA | 7/20/19 | urine | non | H2 |
| KPNMA214 | *K. pneumoniae* | KPC-161 | F | NA | 5/5/22 | urine | non | H3 |
| KPNMA215 | *K. pneumoniae* | KPC-162 | F | 61 | 5/31/22 | rectal swab | CTX-M-15 | H4 |
| KPNMA216 | *K. pneumoniae* | KPC-163 | F | 36 | 8/20/22 | rectal swab | non | H5 |
| KPNMA217 | *K. pneumoniae* | KPC-164 | M | 2 | 8/22/23 | blood | CTX-M-15 | H4 |
| KPNMA221 | *K. pneumoniae* | KPC-44 | M | 45 | 8/19/20 | BAL | non | H4 |
| KPNMA225 | *K. pneumoniae* | KPC-80 | F | 64 | 11/22/19 | Abdominal | non | H6 |
| KPNMA228 | *K. pneumoniae* | KPC-97 | M | 67 | 12/11/20 | rectal swab | non | H7 |
| KPNMA213 | *K. pneumoniae* | KPC-160 | M | 45 | 2/18/20 | urine | CTX-M-15 | H6 |
| KPNMA218 | *K. pneumoniae* | KPC-165 | F | NA | 9/22/22 | urine | CTX-M-15 | H8 |
| KPNMA219 | *K. pneumoniae* | KPC-25 | F | 57 | 12/2/22 | blood | non | H9 |
| KPNMA220 | *K. pneumoniae* | KPC-33 | M | 30 | 5/13/21 | rectal swab | non | H4 |
| KPNMA222 | *K. aerogenes* | KPC-57 | F | 29 | 7/7/20 | rectal swab | non | H10 |
| KPNMA224 | *K. pneumoniae* | KPC-73 | M | 73 | 11/19/22 | BAL | CTX-M-15 | H11 |

NA: not available, BAL: bronco alveolar lavage.

The mutations are grouped based on their structural regions and represented by different colors (refer to table 1). Green- highlighted cells correspond to mutations in Loop 240, pink-highlighted cells denote mutations in Loop 270, blue-highlighted cells indicate Omega-loop mutations, and beige-highlighted cells represent combinations of mutations from different regions
