## Supplementary material for "Emerging Resistance to Novel β-Lactam β-Lactamase Inhibitor Combinations in *Klebsiella pneumoniae* bearing KPC Variants": Suplemental Material: Table S3.pdf

**Table S3**. Antimicrobial resistance genes present in KPNMA215, KPNMA216 and KAMA222.

| DRUG CLASS | ANTIMICROBIAL RESISTANCE GENES | KPNMA215 | KPNMA216 | KAMA222 |
|---|---|---|---|---|
| **AMINOGLYCOSIDES** | *aac(3)-IIa_1* | X | | |
| | *aac(3)-Iid* | | | X |
| | *aac(6')-Ib-cr* | X | | X |
| | *aar-3* | | | X |
| | *strA* | X | X | |
| | *strB* | X | X | |
| **β-LACTAMS** | $bla_{CTX-M-15}$ | X | | |
| | $bla_{KPC-162}$ | | X | |
| | $bla_{KPC-57}$ | | | X |
| | $bla_{KPC-161}$ | X | | |
| | $bla_{OXA-1}$ | X | | X |
| | $bla_{TEM-1B}$ | X | | X |
| | $bla_{SHV-28}$ | X | X | |
| **FOSFOMYCIN** | *fosA6* | X | X | X |
| **MACROLIDES** | *mph(A)* | | | X |
| **SULFONAMIDES** | *sul1* | | X | X |
| | *sul2* | X | | |
| **TETRACYCLINE** | *tet(A)* | X | | X |
| | *tet(D)* | | X | |
| **TRIMETHOPRIM** | *dfrA14* | X | | X |
| **QUINOLONE** | *oqxB* | X | X | X |
| | *oqxA* | X | X | X |
| | *qnrB1* | X | | X |
